# Gene expression of macaques infected with malaria species of zoonotic concern

**DOI:** 10.1101/2023.01.19.524806

**Authors:** Amber E. Trujillo, Christina M. Bergey

**Affiliations:** New York University, Department of Anthropology, New York, NY, USA; New York Consortium in Evolutionary Primatology, New York, NY, USA; Rutgers University, Department of Genetics, New Jersey, USA; Rutgers University, Human Genetics Institute of New Jersey, New Jersey, USA

## Abstract

A multitude of malaria species (genus *Plasmodium*) infects primates. Due to their public health importance, the human-infective species have garnered the most focus, but increased knowledge of non-human primate malaria species is warranted to improve our evolutionary understanding of host-parasite interactions. Additionally, the broad host tropism of some primate malaria parasites and their realized or theorized zoonotic potential add urgency to understanding of primate-parasite interactions. Here, we use comparative transcriptomics to understand the rhesus macaque (*Macaca mulatta*) response to two malaria parasites used as analogues to human-infective species of differing severity and which may represent emerging zoonotic threats: *P. coatneyi*, comparable to human-infective *P. falciparum*, and *P. cynomolgi*, comparable to human-infective *P. vivax*. We first validate our transcriptomics-based proxy of parasite load through comparison to gold-standard microscopy-based measures. We then find that malaria-associated host genes have functional links to immune system regulation and blood cells. Host genes with differing expression by malaria species were more likely to be involved in brain-linked functions, perhaps due to the differential central nervous system involvement of the two parasite species. Such comparative work on primate malaria species may help elucidate the essential and species-specific molecular mechanisms that underlie differing clinical presentations and zoonotic risk.

## Introduction

The five species of malaria that regularly infect humans vary in their clinical presentation and severity, with the most clinically serious being *Plasmodium falciparum* and *P. vivax*. These human-infective parasites are part of lineages that separately came to infect humans thousands of years ago^1^, each having distinct nearest non-human primate-infective relatives. The broad host tropism of *Plasmodium* species is underscored by the recent zoonotic exchange to humans observed for simian malaria species, including the now-widespread *P. knowlesi* ^2, 3^, all of which tend to cause less severe disease than human infection specialist species of *Plasmodium*. However, despite their public health importance, the reasons for the variation in impact between the malaria parasite species remain incompletely understood. Evolutionary transcriptomic investigations of host-parasite interaction can shed light on the molecular underpinnings of this differential pathology^4^. Additionally, understanding the determinants of disease severity has critical relevance for zoonotic disease, as reservoir hosts with infection benign enough to not impact behavior pose the greatest risk of spillover^5^.

Various primate models of malaria, including these zoonotic parasites, have been used to interrogate malaria biology and better understand the host-parasite interaction differences that determine disease severity (e.g., rhesus macaques, *Macaca mulatta*^4, 6–13^; crab-eating macaques, *Macaca fascicularis*^8, 14, 15^; owl monkeys, *Aotus nancymaae*^16^; and squirrel monkeys, *Saimiri boliviensis*^17, 18^). For example, rhesus macaque monkeys infected with *P. coatneyi* are clinically comparable to human infections with the distantly-related *P. falciparum*. Experimental infections in rhesus monkeys cause high virulence with parasitemia often rising rapidly to 10-20%. Symptoms include renal dysfunction, severe anemia, metabolic acidosis, and 30-40% mortality in the absence of effective antimalarial drug treatment^19, 20^. This severity has led to the use of the *P. coatneyi*-rhesus model for *P. falciparum*, particularly for the neurological presentations which may include coma with, importantly, sequestration of infected red blood cells in the cerebral and cerebellar brain vessels as seen in human *P. falciparum* infections^19^. In contrast, when rhesus macaques are instead infected with *P. cynomolgi*, most clinically comparable and also most closely related to human *P. vivax*, the monkeys exhibit a milder disease, which may include acute anemia and thrombocytopenia^21^. Like the human *P. vivax* infections it emulates, mortality risk is low except for pregnant rhesus macaques and fetuses^19^. The utility of the model is also augmented by a *vivax*-like rate of relapses originating from hypnozoites^9, 21–23^.

The host tropism of these parasites provides important context for their differential symptoms in various species and informs their risk for zoonotic spillover. Natural infection with *P. coatneyi* has been observed for several primate species in Southeast Asia, but crab-eating macaques (*M. fascicularis*) are thought to have co-evolved with the parasite^20, 24, 25^. Although pathogenicity in crab-eating macaques is less consistent than in rhesus macaques, these monkeys still variably develop severe symptoms and clinical presentation in response to *P. coatneyi* ^20^. In contrast, *P. cynomolgi* has been found to naturally infect a range of primate hosts in Southeast Asia, including various macaque species^26^. Alongside the established simian zoonotic *P. knowlesi*, both *P. cynomolgi* ^4, 27–30^ and *P. coatneyi* ^30^ parasites were recently found in blood samples of humans, though the latter appears to be far rarer, as supported by prior failed attempts to experimentally infect people^31^. Comparison of host response to these parasites may provide insight into their differing severity and risk of spillover.

Several transcriptomic studies have advanced our understanding of malaria biology by investigating non-human primate response of a macaque model to these two parasites, *P. cynomolgi* and *P. coatneyi* ^4, 32^. Recent studies of *P. cynomolgi*, for example, have uncovered the causes of malaria anemia by examining bone marrow gene expression^10^; determined that genes related to PAMP and pro-inflammatory cytokines were upregulated in severe disease^33^; modeled metabolic pathway systems to determine that patterns of purine metabolism within purine pathways for *P. cynomolgi* infection were consistent with *P. falciparum* infection in humans^11^; and found many more genes were upregulated in acute vs. relapsing disease with enrichment for involvement in immune response^12^. Transcriptomic investigations of macaque response to *P. coatneyi* are more limited, but extensive systems biology data exist from controlled experiments^22^. Such studies of experimentally-infected macaques have greatly improved our understanding of fundamental malaria biology^4, 32^. However, to date, no direct comparison of the gene expression response to *P. coatneyi* versus *P. cynomolgi* has been undertaken, limiting our understanding of how host response to the parasites may contribute to the differential disease severity and signs.

In this study, we compare two cohorts of rhesus macaques (*Macaca mulatta*) that have been experimentally infected with either *P. cynomolgi* and *P. coatneyi* for longitudinal analysis. We partition the macaque gene expression response to infection into components that are generalized or specific to infecting *Plasmodium* species, thereby identifying fundamental processes to malaria in this host as well as changes which may underpin the unique symptoms of each species. We also simultaneously assay *Plasmodium* gene response, determining orthologous genes with similar or divergent reactions across the two parasite species. Overall, our comparison of macaque interaction with *P. cynomolgi* or *P. coatneyi* allows insight into the transcriptomic drivers of primate-parasite interaction in general, as well as the use of these systems to model human malaria with different parasite etiology.

## Results

The rhesus macaque whole blood transcriptomic datasets were generated through the efforts of the Malaria Host-Pathogen Interaction Center’s (MaHPIC) to test hypotheses related to host-pathogen interactions in experimental non-human primate malaria infections. The *N* = 50 samples were from eight macaque individuals from a captive, experimental population. To identify macaque monkey genes with expression correlated to *Plasmodium* parasite levels, we aligned blood transcriptome reads to a concatenated host-parasite pseudo-reference genome (containing genomes of macaque and *Plasmodium coatneyi* or macaque and *P. cynomolgi*) and tallied reads by exon. After filtering, the median per-sample counts of reads uniquely mapping to the macaque and *P. coatneyi* genome were 30,987,028 (range: 9,945,658 to 75,084,213) and 24,177 (range: 2,899 to 26,416,731), respectively. For the macaque and *P. cynomolgi* dataset, the median uniquely mapping per-sample counts were 38,758,474 (range: 3,071,635 to 99,902,110) and 28,793 (range: 7,017 to 29,629,054), respectively.

### Inferred parasitemia proxy is an accurate representation of infection

To investigate the accuracy of the parasite load proxy inferred computationally from RNA sequencing data, we tested for a correlation between these inferred parasite load proxies and previously published parasite load counts measured via microscopy as parasites per microliter, a method commonly used. We found that our parasite load proxy: *i.)* was tightly and significantly correlated with the traditional measure of parasite load (Fig. 1a; R^2^=0.8527, *p* < 2.2 × 10^−16^); and *ii.)* accurately traced the severity of parasite load over the 100-day experimental period with high sensitivity (Fig. 1b). Having found this proxy to be a reliable method for inferring parasite load from RNAseq data, later analyses used this metric.

**Figure 1.**
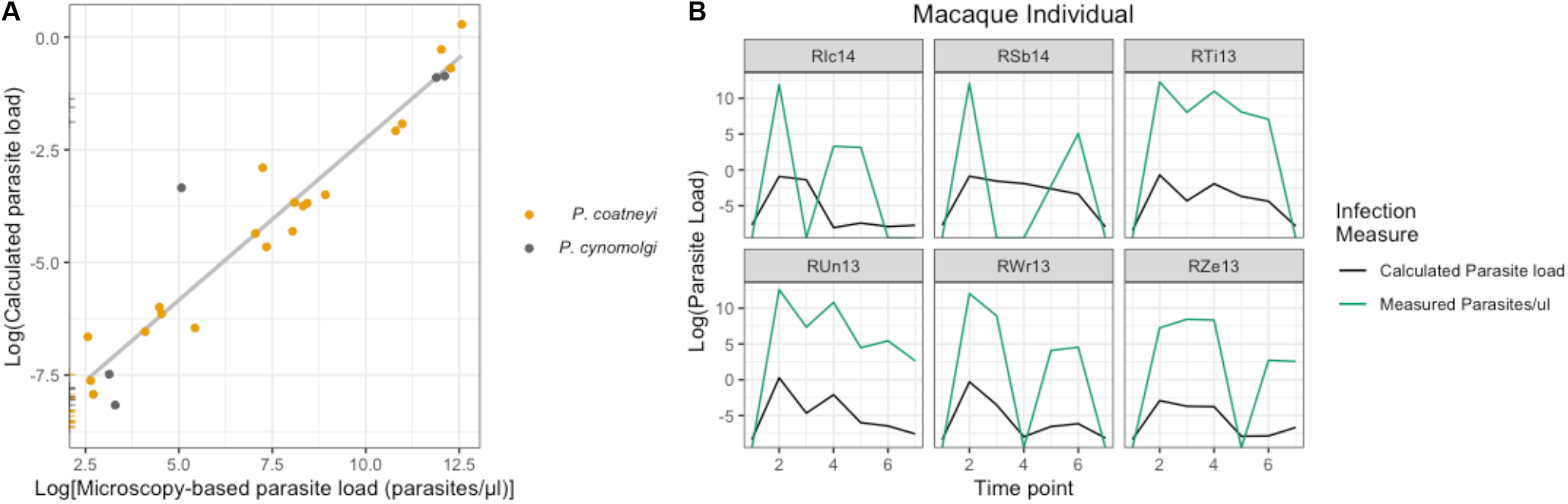
Correlation between previously published microscopy-based parasite load and the sequence read-based parasite load for samples infected with *P. coatneyi* (yellow) and *P. cynomolgi* (black). Samples were considered to be uninfected if there was no evidence of infection via microscopy and the proxy value was less than 0.001. Lines indicate samples with microscopy values of 0. Measures of infection were significantly and highly correlated (*R*^2^ = 0.853, *p* < 2.2 × 10^−16^; **A**) each capturing parasite load changes throughout the 100 day experiment (**B**). Samples from individuals RCs13 and RFv13 are excluded from the plot given their lack of malaria infection and removal from the experiment, respectively.

### Contrasting host response to parasitemia and infection intensity

In order to explore how the host may be responding to the malaria parasite, we analyzed our dataset at two hierarchical levels: parasitemia (infected vs. non-infected) and parasite load (inferred intensity of infection.) In analyzing *N* = 50 longitudinal samples from five individual macaques infected with *Plasmodium coatneyi* and from three individuals infected with *P. cynomolgi*, we found that there were significant differences in the identified genes of interest between the two measures of infection with no overlap between the top 20 DE genes (Table 1, Table 2). Nevertheless, DE genes between the two measures were all associated with similar functions and disorders pertaining to immune system regulation and blood cells. For example, when considering parasite load (Fig. 2a), significantly DE genes include *ADAMDEC1* (LogFC=3.67, adj. p=2.66 × 10^−10^), *CYP2C76* (LogFC=2.35, adj. p=6.45 × 10^−7^), *SLCO2B1*, (LogFC=3.70, adj. p=7.00 × 10^−7^), *CA4* (LogFC=4.25, adj. p=1.90 × 10^−6^), *RABGAP1L* (LogFC=−0.604, adj. p=1.76 × 10^−4^), and *LUC7L* (LogFC=−0.517, adj. p=0.007). When considering parasitemia (Fig. 2b), significantly differentially expressed genes include *GAS6* (LogFC=2.08, adj. p=4.95 × 10^−5^) and *PTPN1* (LogFC=0.609, adj. p=4.95 × 10^−5^). Results for DE genes identified using the coefficients of the *Plasmodium* species and interacting variables (infection measure\**Plasmodium* species) for each model are given in the supplementary tables, as are full results for parasite load- and parasitemia-DE tests (Tables S1-S2).

**Table 1.** Macaque genes with the strongest evidence of differential expression by malaria parasitemia (infected versus non-infected samples), irrespective of *Plasmodium* species. Table is limited to 20 genes, with full results present in a supplementary table (Tables S1).

| Macaque genes with expression DE for parasitemia |  |  |  |  |  |
| --- | --- | --- | --- | --- | --- |
| Positively correlated |  |  | Negatively correlated |  |  |
| Gene | *logFC* | Adj. *p* | Gene | *logFC* | Adj. *p* |
| *HIST1H2BH* | 1.96 | $2.82 \times 10^{-5}$ | *APBB2* | -1.09 | $4.97 \times 10^{-5}$ |
| *GAS6* | 2.08 | $4.95 \times 10^{-5}$ | *DHRS3* | -0.975 | $7.81 \times 10^{-5}$ |
| *PTPN1* | 0.609 | $4.95 \times 10^{-5}$ | *PDE3B* | -0.645 | $2.15 \times 10^{-4}$ |
| *ELL3* | 1.40 | $4.95 \times 10^{-5}$ | *NXPH3* | -1.68 | $2.47 \times 10^{-4}$ |
| *SOCS1* | 1.34 | $4.95 \times 10^{-5}$ | *CUX2* | -1.61 | $2.70 \times 10^{-4}$ |
| *ASAP3* | 1.18 | $7.81 \times 10^{-5}$ | *MAPT* | -0.834 | $2.96 \times 10^{-4}$ |
| *AGRN* | 1.45 | 0.000114 | *PID1* | -1.73 | $2.99 \times 10^{-4}$ |
| *FADS1* | 1.25 | 0.000158 | *ESRP2* | -0.564 | $2.99 \times 10^{-4}$ |
| *TSTD1* | 0.813 | 0.000172 | *VANGL1* | -0.932 | $2.99 \times 10^{-4}$ |
| *EGR2* | 1.34 | 0.000247 | *CA8* | -1.06 | $3.10 \times 10^{-4}$ |
| *PIM2* | 0.609 | 0.000247 | *NFATC3* | -0.569 | $3.10 \times 10^{-4}$ |
| *ANKRD9* | 1.85 | 0.000247 | *XPC* | -0.540 | $3.10 \times 10^{-4}$ |
| *RELB* | 0.60 | 0.000247 | *DBP* | -0.663 | $3.10 \times 10^{-4}$ |
| *AICDA* | 2.66 | 0.000247 | *AOX2* | -1.11 | $3.18 \times 10^{-4}$ |
| *ZNF215* | 0.779 | 0.000247 | *CD1B* | -2.14 | $4.25 \times 10^{-4}$ |
| *FADS2* | 2.32 | 0.000247 | *PIK3R1* | -0.468 | $4.25 \times 10^{-4}$ |
| *FFAR1* | 2.09 | 0.000247 | *SAMD3* | -0.941 | $6.60 \times 10^{-4}$ |
| *GPC1* | 2.49 | 0.000247 | *C1H1orf174* | -0.388 | $6.62 \times 10^{-4}$ |
| *P2RX5* | 1.31 | 0.000270 | *KLRB1* | -1.33 | $7.57 \times 10^{-4}$ |
| *JAK3* | 0.675 | 0.000270 | *CD96* | -0.731 | $7.64 \times 10^{-4}$ |

**Table 2.** Macaque genes with the strongest evidence of differential expression by malaria parasite load (amount of parasite RNA detected in sample), irrespective of *Plasmodium* species. Table is limited to 20 genes, with full results present in a supplementary table (Table S2).

| Macaque genes with expression DE for parasite load |  |  |  |  |  |
| --- | --- | --- | --- | --- | --- |
| Positively correlated |  |  | Negatively correlated |  |  |
| Gene | *logFC* | Adj. *p* | Gene | *logFC* | Adj. *p* |
| *ADAMDEC1* | 3.67 | $2.66 \times 10^{-10}$ | *ASPH* | -1.19 | $7.68 \times 10^{-6}$ |
| *FAM20A* | 2.76 | $2.66 \times 10^{-10}$ | *CAMK1D* | -0.724 | $1.82 \times 10^{-5}$ |
| *KCNE1* | 4.02 | $1.18 \times 10^{-7}$ | *RABGAP1L* | -0.604 | $1.76 \times 10^{-4}$ |
| *MARCO* | 4.02 | $6.01 \times 10^{-7}$ | *ADGRG5* | -1.54 | $4.40 \times 10^{-4}$ |
| *CYP2C76* | 2.35 | $6.45 \times 10^{-7}$ | *PIK3R6* | -1.92 | $5.41 \times 10^{-4}$ |
| *PLXDC1* | 2.85 | $7.00 \times 10^{-7}$ | *PDE7A* | -0.959 | $6.66 \times 10^{-4}$ |
| *SLCO2B1* | 3.71 | $7.00 \times 10^{-7}$ | *MAMU.DMB* | -1.19 | $7.95 \times 10^{-4}$ |
| *PODXL* | 1.93 | $7.00 \times 10^{-7}$ | *ADCY9* | -1.27 | $7.96 \times 10^{-4}$ |
| *HSD3B7* | 2.10 | $7.00 \times 10^{-7}$ | *ADK* | -1.34 | $9.82 \times 10^{-4}$ |
| *CDH17* | 3.31 | $7.87 \times 10^{-7}$ | *EEF2K* | -0.744 | $9.87 \times 10^{-4}$ |
| *CTH* | 4.32 | $7.87 \times 10^{-7}$ | *TXNRD2* | -0.860 | $1.44 \times 10^{-3}$ |
| *FCGR1A* | 3.47 | $1.19 \times 10^{-6}$ | *ABCC4* | -1.31 | $1.45 \times 10^{-3}$ |
| *LIPM* | 2.46 | $1.23 \times 10^{-6}$ | *ABAT* | -0.981 | $1.49 \times 10^{-3}$ |
| *TIFA* | 2.54 | $1.30 \times 10^{-6}$ | *DNAJB12* | -0.428 | $1.67 \times 10^{-3}$ |
| *SYNPO2* | 2.44 | $1.48 \times 10^{-6}$ | *ULK2* | -0.865 | $1.94 \times 10^{-3}$ |
| *NLRP14* | 3.08 | $1.54 \times 10^{-6}$ | *BTAF1* | -0.758 | $2.41 \times 10^{-3}$ |
| *TIMP3* | 3.21 | $1.54 \times 10^{-6}$ | *SLC4A7* | -0.652 | $2.45 \times 10^{-3}$ |
| *P2RY14* | 2.97 | $1.54 \times 10^{-6}$ | *MAMU.DMA* | -1.26 | $2.61 \times 10^{-3}$ |
| *TUSC3* | 3.31 | $1.54 \times 10^{-6}$ | *EZH1* | -0.525 | $3.36 \times 10^{-3}$ |
| *NCKAP1* | 3.11 | $1.58 \times 10^{-6}$ | *ADRB2* | -1.38 | $3.68 \times 10^{-3}$ |

**Figure 2.**
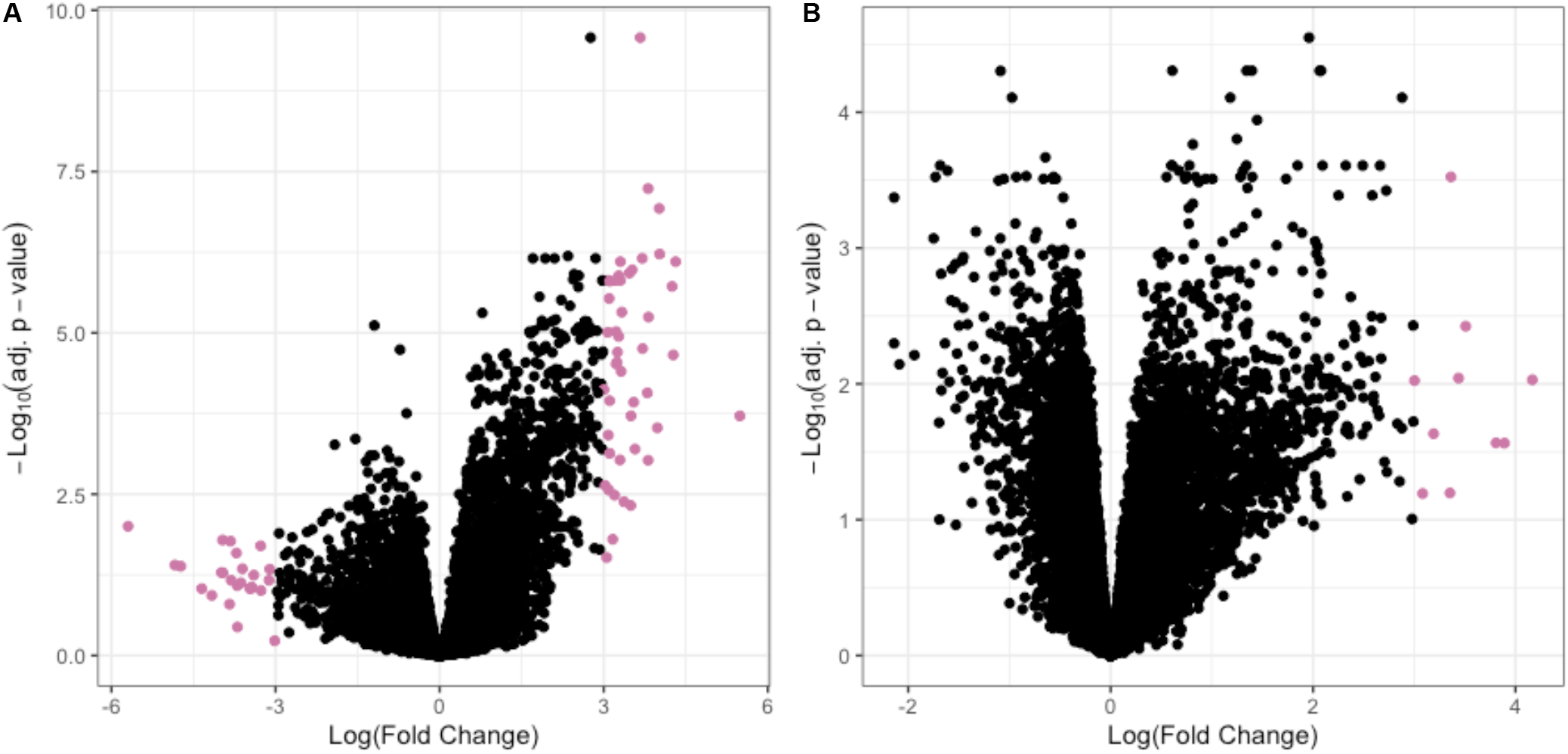
Summary of tests for differential expression of macaque genes by malaria infection status. Pink dots indicate differentially expressed macaque genes for two different measures of malaria infection: parasite load, or the inferred intensity of infection (**A**), and parasitemia, or samples with and without detected malaria infection (**B**). Both measures are inferred from sequencing reads. Note differing scales between two graphs.

To further investigate the impact of the various parameters on individuals’ gene expression, we conducted Principle Components Analysis (PCA; Fig. 3a and b) using the combined dataset of macaques infected with different species of *Plasmodium* and considering the two measures of infection. We found that individuals were most strongly grouping according to their parasitemia status (uninfected or infected) (PC1; Fig. 3a) with secondary grouping according to the infecting *Plasmodium* species (PC3; Fig. 3b). There was no distinct grouping according to parasite load. We additionally performed t-Distributed Stochastic Neighbor Embedding (t-SNE) analysis, but found no significant clustering.

**Figure 3.**
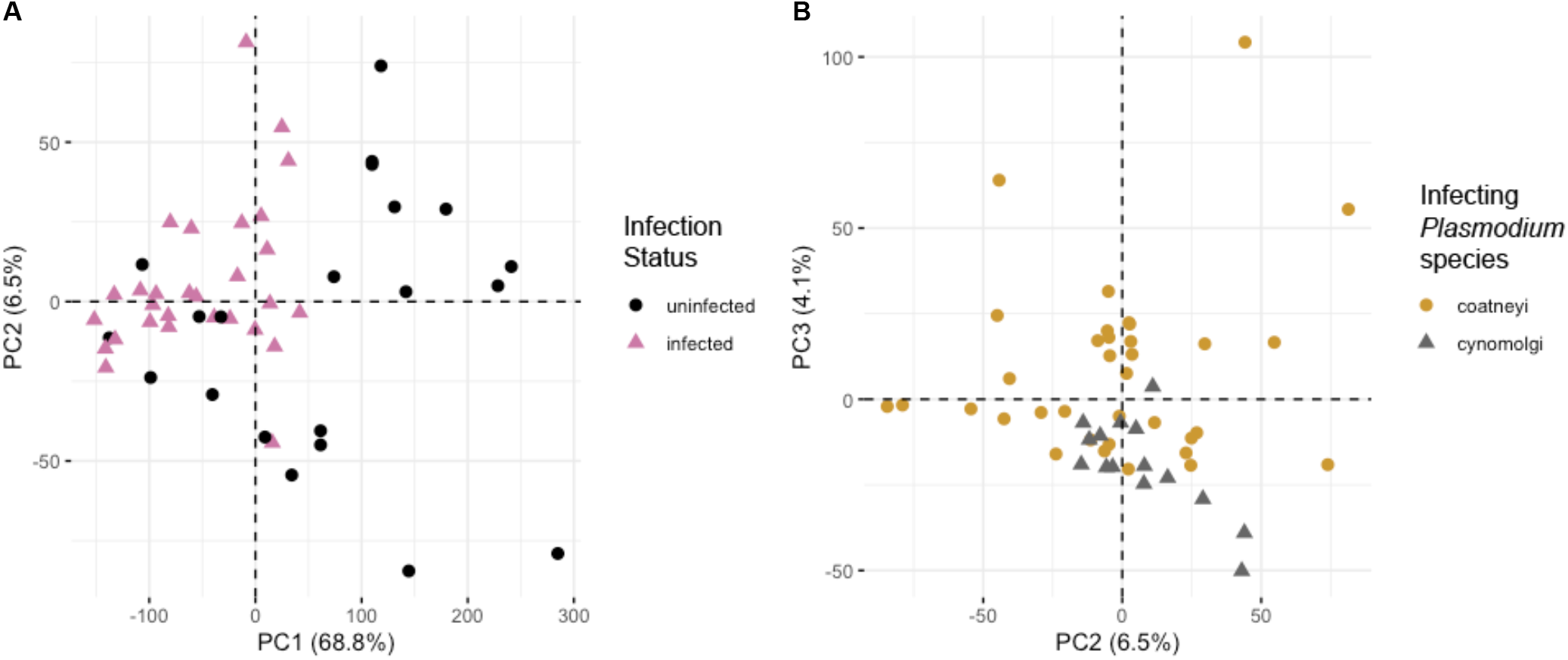
Visualization of Principal Components Analysis (PCA) of macaque gene expression. (**A**) The first principal component, explaining 68.8% of the variation, corresponded approximately to parasitemia, distinguishing infected (pink triangles) versus non-infected (black circles) samples. (**B**) The third principal component, explaining 4.1% of the variation, partially distinguished samples infected with differing *Plasmodium* species: *P. coatneyi* (yellow circles) versus *P. cynomolgi* (gray triangles).

### Over-represented functions of macaque genes correlated to malaria infection

Finally, we found that up-regulated genes DE for both of the measures of parasite infection (parasite load and parasitemia) were associated with functions such as immune system process (GO:0002376; adj. p=2.719 × 10^−35^ and 1.93 × 10^−8^, respectively), response to stimulus (GO:0050896; adj. p=6.54 × 10^−32^ and 1.95 × 10^−15^, respectively), and response to stress (GO:0006950; adj. p=7.09 × 10^−30^ and 1.74 × 10^−6^, respectively). We found that DE genes that were detected using parasite load specifically were enriched for functions such as defense response (GO:0006952; adj. p=6.68 × 10^−27^), response to other organisms (GO:0051707; adj. p=4.41 × 10^−18^), and platelet activation (KEGG:04611; adj. p=2.803 × 10^−4^), while DE genes that were detected using parasitemia were enriched for functions such as increased red cell osmotic fragility (HP:0005502; adj. p=2.74 × 10^−7^), anemia due to reduced life span of red cells (HP:0011895; adj. p=1.88 × 10^−5^), and reticulocytosis (HP:0001923; adj. p=2.74 × 10^−7^).

When exploring the DE-genes associated with the infecting *Plasmodium* species coefficient using parasitemia, we found that *P. coatneyi* (which is comparable to the *Plasmodium* species that causes cerebral malaria in humans) genes were particularly enriched for vestibular hypofunction (HP:0001756; adj. p=1.716 × 10^−2^), abnormal cochlea morphology (HP:0000375; adj. p=3.021 × 10^−2^), and subcortical cerebral atrophy (HP:0012157; adj. p=3.021 × 10^−2^; Table S3). There were no unique functional enrichment categories of interest to report when including the DE-genes associated with the infecting *Plasmodium* species coefficient using parasite load (Table S4). We found that none of our differentially expressed gene lists that were detected using the *Plasmodium* species coefficient (*i.e.*, *P. coatneyi*- or *P. cynomolgi*-specific genes) were more than likely by chance to be overrepresented in human response genes to *Plasmodium* parasites (i.e., *P. vivax* and *P. falciparum*^34^) with the exception of genes upregulated in *P. cynomolgi* relative to *P. coatneyi* , as these genes were more likely than expect by chance to be upregulated in *P. falciparum* response (p value=0.034).

Results for the functional enrichment analyses of macaque genes DE for the interaction between infection measure and *Plasmodium* species are included in the supplementary tables (Tables S3-S4).

### Malaria parasite response across infection

To explore further the relationship between the host and pathogen during infection, we identified orthologous genes of the two *Plasmodium* species (Tables S5-S6) and then tested for those with expression correlated to parasite load, the infecting *Plasmodium* species, or the interaction between parasite load and *Plasmodium* species (Table S7). Here, we report the *P. coatneyi* gene accession for the orthologous pairs. For the parasite load variable we found no significantly up-regulated genes but 151 down-regulated *Plasmodium* genes including PCOAH 00027310 (LogFC= −7.224; adj. p=1.36 × 10^−5^), which encodes an ATPase and has been shown to be essential for invasion, as *Plasmodium* uses ATP to determine which host cell to invade (Gunalan et al., 2013) and PCOAH 00031020 (LogFC=−6.947; adj. p=3.66 × 10^−5^), which encodes a highly conserved protease that is involved in the release of merozoite from erythrocyte and cell invasion (Withers-Martinez et al., 2012). For the *Plasmodium* species predictor, we found 351 *P. cynomolgi*-associated genes (*e.g.*, PCOAH 00025970: LogFC=4.722; adj. p=3.28 × 10^−13^) and 310 *P. coatneyi*-associated genes (*e.g.*, PCOAH 00034280: LogFC=−5.034; adj. p=1.76 × 10^−14^). Finally, for the interaction variable (parasite load\**Plasmodium* species) we found no significantly up-regulated genes and only 20 down-regulated genes (*e.g.*, PCOAH 00021590: LogFC=−13.480; adj. p=6.133 × 10^−4^); and PCOAH 00041960: LogFC=−17.955; adj. p=6.05 × 10^−5^).

In order to further explore the relationships between gene expression and biological factors, we conducted PCA and t-SNE analyses. The PCA clustered samples according to the degree of parasite load (PC1) and infecting *Plasmodium* species (PC2) (Fig. 4a). The t-SNE analysis suggested that *P. coatneyi* expression was most similar to that of *P. cynomolgi* when parasite load was high (Fig. 4b).

**Figure 4.**
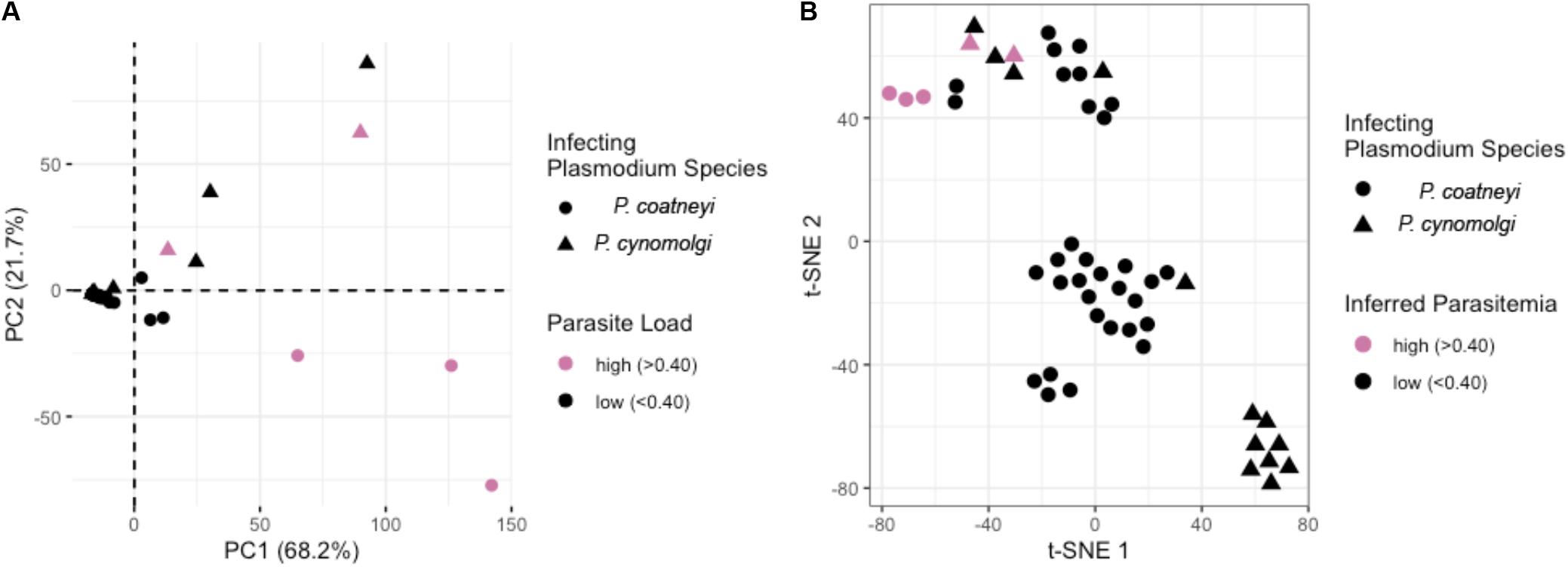
(**A**) Visualization of Principal Components Analysis (PCA) of *Plasmodium* gene expression. The first principal component, explaining 68.2% of the variation, captured parasite load, distinguishing high (pink) versus low (black) parasite load samples. The second principal component, explaining 21.7% of the variation, distinguished samples infected with differing *Plasmodium* species: *P. coatneyi* (circles) versus *P. cynomolgi* (triangles). (**B**) Visualization of results of t-Distributed Stochastic Neighbor Embedding (t-SNE) analyses on macaque gene expression with perplexity of nine. Samples clustered largely by infecting *Plasmodium* species: *P. coatneyi* (circles) versus *P. cynomolgi* (triangles). Of note, *P. coatneyi*-infected macaque samples with the highest parasite load (pink circles) clustered with *P. coatneyi*.

### Over-represented functions of malaria genes linked to parasite load

We conducted functional enrichment analyses using annotations from PlasmoDB (Bahl et al. 2003) for the orthologous gene pairs in *P. coatneyi* and *P. cynomolgi* that were significantly differentially expressed for parasite load, *Plasmodium* species, or their interaction (Table S8). Parasite load-DE genes were enriched for reproduction (GO:0000003; p-value=0.003) and microtubule-based movement (GO:0007018; p-value=5.51 × 10^−5^). For the *Plasmodium* species-DE genes, we found that *P. cynomolgi*-associated genes were enriched for multiple binding molecular functions including ion binding (GO:0043167; p-value=6.96 × 10^−6^) and microtubule binding (GO:0008017; p-value=0.009), and *P. coatneyi*-associated genes were enriched for transferase activities including those involved in one-carbon (GO:0016741; p-value=0.030) and nitrogenous groups (GO:0016769; p-value=0.0466). Finally, for genes DE for the interaction of *Plasmodium* species and parasite load, *i.e.*, those that respond differentially to high parasite load between species, these DE *Plasmodium* genes were enriched for integral (GO:0016021; p-value=0.017) and intrinsic (GO:0031224; p-value=0.017) components of the membrane.

## Discussion

Primates (including humans) and malaria have been co-evolving for thousands of years, leading to the evolution of both shared and species-specific coping strategies in both host and parasite. These primate-parasite relationships have created an opportunity to study model non-human primates in order to understand the biological mechanisms that are involved in human infections that differ in severity and symptomatology. For example, macaques infected with *Plasmodium coatneyi* serve as a useful model for human *P. falciparum* infections, particularly for the study of cerebral malaria^35–37^, and for malaria during pregnancy^20^. In contrast, macaques infected with *P. cynomolgi* have been used to model human *P. vivax* infections, particularly relapsing malaria in which both *Plasmodium* species can remain dormant in the liver as hypnozoites^4^. The differential symptoms are also important factors in determining the risk of zoonotic spillover to humans, as infections that are sufficiently benign to not impact behavior may be hypothesized to pose a greater spillover risk. This is in accordance with observations for these macaque malarias, as spillover appears rare for the more dangerous *P. coatneyi* but increasingly observed for the lower severity *P. cynomolgi*. This differential severity and spillover risk motivated our comparison of macaque host response to these two parasites.

As has been previously established for human patients infected with *P. falciparum* (e.g.,^38^), we confirmed that the proportion of whole blood RNA sequencing reads that map to the *Plasmodium* genome can be successfully used as a proxy for both parasitemia (presence or absence of parasite) and parasite load (proportion of infected red blood cells in macaques infected with *P. coatneyi* and *P. cynomolgi*). Owing to the release of microscopy data by the MaHPIC project, we were able to validate this measure in macaques to demonstrate that our bioinformatic proxy: *i.)* was tightly correlated with the traditionally-used microscopic measure of parasitemia/parasite load, and *ii.)* mirrored parasite load levels over the infection period. These findings support the use of this measure in future studies of wild populations for which microscopic measures of parasite load are infeasible. As sequencing data generation becomes increasingly inexpensive, bioinformatic inference of parasitemia or parasite load may make this computational method preferable to labor intensive microscopy.

Across the two *Plasmodium* species, we identified many significantly differentially expressed genes in the macaque host correlated with parasite load and parasitemia associated with immune function, red blood cell function, and morphology. Many of these genes were similarly linked to malaria infection in humans, particularly those we identified as correlated with the magnitude of parasite load or parasitemia. Such up-regulated genes include *ADAMDEC1* which is involved in inflammatory response during placental *P. falciparum* malaria^39^, *SLCO2B1* which has been associated with parasitemia clearance rate in human *P. vivax* infection^40^, and *CA4* which is negatively correlated with hemoglobin concentrations in children with *P. falciparum* malaria^41^. Similarly down-regulated genes were often implicated in human malaria response. These include *RABGAP1L* which has roles in hematopoiesis^42^ and has been found to have been under selection^43^; *LUC7L* which contains SNPs associated with malaria prevalence in a Chinese population and with the blood disorder *α*-thalassemia that is protective against malaria^44^; and *ADRB2* which contains SNPs significantly associated with malaria in a Southern Indian population^45^. Such overlap between rhesus macaque genes responsive to malaria and human genes linked to malaria susceptibility emphasize the similarity across these two primates in response to diverse malaria parasites and supports the suitability of these non-human primate models for research aiming to improve our fundamental understanding of malaria.

The parallel infection of rhesus macaques with *Plasmodium coatneyi* and *P. cynomolgi* allowed us comparative insight into malaria species-level differences that may mirror those seen between human *P. falciparum* and *P. vivax* infections. Although both macaque malarias are more closely related to *P. vivax*, *P. coatneyi* infection of macaques results in *falciparum*-like cerebral malaria and associated high mortality. Consistent with this severity, genes involved in the macaque response to *P. coatneyi* specifically (in contrast to *P. cynomolgi*) were significantly enriched for relationships to brain morphology and function such as subcortical cerebral atrophy. No phenotypes were enriched for *P. cynomolgi* species-specific response genes, in contrast. This association between *P. coatneyi* and brain-associated functions indicates that this *Plasmodium* species may be inducing transcriptomic response that is similar to the comparable *P. falciparum* infection in humans, supporting use of this model in studies of cerebral malaria.

There are several noteworthy limitations to our study, which was possible only through the generous data sharing of the original MaHPIC project. The most important caveat is that *Plasmodium* species and experimental trial were conflated in our analysis. Although some of the impact of technical variation can be mitigated during bioinformatic library normalization, it is likely that some differentially express genes which we attribute to malaria species-specific response are instead due to differences in experimental setup, macaque genetics, or biases introduced during RNA extraction, cDNA synthesis, library preparation, or sequencing. An improved experimental design to identify malaria species-specific response could be structured to minimize these confounding factors in the future. However, our prior hypothesis of *falciparum*-like nervous system involvement in *P. coatneyi* infections lends support to our finding of a broad enrichment of brain-associated functions, which is unlikely attributable to technical artifacts.

As treatment regimens varied between the studies, were subcurative, and were timed to coincide with high parasite load, several genes and pathways we identified as being differentially expressed in response to malaria may instead be linked to anti-malaria drug response. For instance, we found *CYP2C76* to be upregulated in response to parasite load which has previously been attributed to metabolism of the anti-malarial mefloquine in crab eating macaques (*M. fascicularis*)^46^. Additionally, genes that were differentially responsive to *P. coatneyi* parasitemia were enriched for vestibular hypofunction and abnormal cochlear morphology, which may reflect the acute ototoxicity commonly associated with certain antimalarials^47, 48^. Here again, malaria species and antimalarial drug are confounded: The *P. coatneyi*-infected macaques were given subcurative and curative treatments of artemether alone while the *P. cynomolgi*-infected macaques were given subcurative blood-stage treatment of artemether and curative treatment of chloroquine and primaquine. Quinoline derivatives such as chloroquine and primaquine are known to demonstrate ototoxicity, but the risk for artemisinin derivatives like artemether remains controversial^47^. Future study of macaque transcriptomic response to antimalarials may support our findings and could lead to insight into the pathways involved in ototoxicity and identification of biomarkers for this important side-effect.

Overall, our investigation into the gene expression response of a primate model to two malaria parasites of emerging concern identified many similarities with the comparatively well-studied human malaria response. This common architecture of *Plasmodium* transcriptomic response supports the use of experimentally-infected macaques to better understand fundamental primate malaria pathophysiology to improve human treatment. Similarly our comparative study revealed brain-associated differences between the macaque response to malaria species *P. coatneyi* and *P. cynomolgi*, supporting the use of the former in models of *P. falciparum* cerebral malaria. Future comparative studies of non-human primate response to malaria and malaria-like parasites can benefit from the sequencing read-based proxy, which we newly validated in these species. Such “dual-transcriptomic” research that simultaneously assays host and parasite response will allow insight into primate-parasite coevolution and may identify new therapeutic targets for human malaria. Finally, there is pressing practical motivations to study these parasites in particular: Although *P. coatneyi* and *cynomolgi* are traditionally deemed simian malaria species, their recent identification in human blood samples heightens the necessity of understanding their interaction with their primate hosts, both macaque and human.

## Methods

### Experimental set-up and dataset

We analyzed RNA sequence (mRNA-seq) data previously generated by the Malaria Host-Pathogen Interaction Center (MaHPIC)^6, 22^. The datasets used were generated from 50 whole blood samples of 5 malaria-naïve rhesus macaque (*Macaca mulatta*) individuals that had been experimentally inoculated with *Plasmodium coatneyi* Hackeri strain (study PRJNA400695) and 5 malaria-naïve rhesus macaque individuals that had been experimentally inoculated with *Plasmodium cynomolgi* strain B (study PRJNA388645). Methods for sample collection, pathogen inoculation, RNA extraction, library preparation, and transcriptome sequencing are fully described in the Gene Expression Omnibus page (GSE103259 and GSE99486, respectively) for these experiments.

Briefly, for the first experiment, researchers in the MaHPIC consortium^6^ infected 5 adult malaria-naïve male macaques with *P. coatneyi* sporozoites from the salivary glands of *Anopheles dirus*, *An. gambiae*, and *An. stephensi* mosquitoes, which were produced and isolated at the Centers for Disease Control and Prevention. These macaques were then monitored for 101-days. Subcurative treatment of artemether was administered to all subjects at the initial peak of infection and complete blood-treatment with artemether was given at the end of the experiment. Only one individual (RCs13) did not develop infection and was not given additional subcurative treatments for subsequent recrudescence peaks. Whole blood samples were taken at 7 time-points over the 101-day period (35 samples total). Parasite load for each time point was measured as follows: 1) pre-infection or uninfected control; 2) acute infection or the peak infection as determined by symptomatic state and parasitological assessment; 3) early post-subcurative treatment or the earliest observation after peak infection and treatment; 4) late post-subcurative treatment; 5) early chronic or observation of low levels of persistent parasitemia; 6) late chronic; and 7) final measurement at end of experimental period.

For the second experiment, researchers in the MaHPIC consortium^22^ infected 5 adult malaria-naïve male macaques with *P. cynomolgi* sporozoites from the salivary glands of *Anopheles dirus*, *An. gambiae*, and *An. stephensi* mosquitoes, which were produced and isolated at the Centers for Disease Control and Prevention. These macaques were also monitored for 101-days and subcurative treatment was administered to all subjects at the initial peak of infection. One individual (RFv13) was euthanized during the experiment due to clinical complications of malaria and only two blood samples were taken. Whole blood samples were taken at 7 time-points over the 101-day period (30 samples total; 29 from the surviving subjects.) RNAseq data was not collected for RSb14 at time point 5, and we exclude a blood sample taken from the transfusion donor that was used for individual RFv13 to treat severe clinical complications. For the present study, individuals that experienced phenotypically severe malaria (RMe14 and RFa14) were also excluded as they were given alternative treatments to cope with clinical complications. Blood was sampled and parasite load was measured across the following time points: 1) pre-infection control; 2) at acute infection or the peak infection as determined by symptomatic state and parasitological assessment; 3) 7 days post-peak infection; 4) during relapses; 5) during inter-relapse intervals; and 6) at the end of the experimental period.

### Gene expression inference

For both RNA-seq datasets we removed Illumina-specific adapter regions (using parameters ILLUMINACLIP:TruSeq3-PE.fa:2:30:10) and low quality bases (quality score < 3) at the beginning and end of each raw sequencing read, and trimmed the read when the average quality of bases in a sliding 4-base wide window fell below 15 using Trimmomatic^49^. All statistical analyses were performed in R^50^ unless otherwise stated. We removed reads that were less than 36 basepairs after trimming. In order to determine if sequence reads were from the host, we performed a competitive mapping analysis as follows. Using the STAR alignment program^51^, we mapped trimmed, quality-filtered reads to concatenated reference genomes that include rhesus macaque (GCF 003339765.1 Mmul 10) and parasite (*P. coatneyi* : GCF 001680005.1 ASM168000v1^52^; or *P. cynomolgi*: GCF 000321355.1 PcynB^53^), allowing each read to map multiple locations with up to eight mismatches per 100 basepair paired-end read (after^54^). The best matching, primary alignments were extracted and then merged into pathogen and host “unique” datasets (*i.e.*, primary reads that have only mapped to the host or parasite), while excluding unplaced scaffolds for macaque.

We used Rsubread^55^ to create a macaque and *Plasmodium* read-count matrix for each host individual, using the “uniquely” mapped host and pathogen reads, respectively, and allowing for overlapping reads. Reads for each analysis were tallied at the level of exons which were then summed to create a per-gene count for the sample, using gene annotations (GCF 003339765.1 Mmul 10 and GCF 001680005.1 ASM168000v1 for the first dataset, and GCF 003339765.1 Mmul 10 and GCF 000321355.1 PcynB for the second dataset).

To compare *Plasmodium* gene expression between the two experiments (the first using *P. coatneyi* and the second, *P. cynomolgi*) from the two macaque cohorts, we combined the above *Plasmodium* read count matrices matched by orthologous genes. We identified orthologous genes using orthologue groups from PlasmoDB^56^ and excluded any genes for which orthologous genes between the two *Plasmodium* species did not exist. This combined read count matrix was then used to explore *Plasmodium* differentially expressed genes responding to parasite load.

### Correlation of sequencing read- and microscopy-based parasite load inference

We inferred a proxy of parasite load (“sequencing read-based parasite load”) for all samples by calculating the ratio of the number of *Plasmodium* pathogen reads to the number of host macaque reads recovered (after^38, 57^). To validate this metric, we tested for a correlation between our parasite load proxy and previously published microscopy-based parasite load calculated from thick and thin blood smears for these samples. Microscopy-based parasite load was defined as parasites count per microliter of whole blood taken on the same day as RNA data was collected. We tested whether our inferred parasitemia proxy and the original studies’ microscopy-based estimates were significantly correlated using a linear regression model comparing our calculated parasite load against the microscopy-based estimate (parasites per microliter) and by computing Spearman’s correlation coefficient.

### Identification of differentially expressed host genes across two measures of pathogen infection and infecting malaria species

We next identified rhesus macaque genes with expression correlated to either parasite load (relative amount of parasite in blood) or parasitemia (presence of parasite in blood), as well as the species of infecting *Plasmodium*. To do so, we estimated the sequencing read-based parasite load (hereafter “parasite load”) as the ratio of pathogen to host reads, and sequencing read-based parasitemia (hereafter “parasitemia”), which we defined as positive if the proxy were greater than 0.001 with confirmation from microscopy. Using the combined read count matrix (with data from PRJNA400695 and PRJNA388645), we first calculated normalization factors and filtered for lowly expressed genes, defined as any gene with expression below a counts per million (CPM) threshold of 1 in at least 1 individual for macaque genes using the Bioconductor package edgeR^58^. With these calculated normalization factors, we then used the limma-voom package^59, 60^ to include biological replicates as a random effect (i.e., blocking variable) and each parasitemia measure (parasite load and parasitemia) as a fixed effect in the following models:

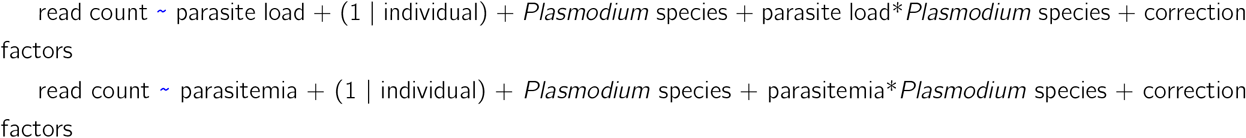

where correction factors include those adjusting for differences in number of reads and library complexity. We extracted the coefficients and adjusted *p*-values for all predictors for each gene from this model, correcting for multiple tests via FDR. We considered a gene to be a parasitemia-DE gene if the adjusted *p*-value was less than 0.05.

To explore potential biological relationships (*e.g.*, between parasite load or parasitemia and infecting *Plasmodium* species) using normalized and filtered read counts, we performed dimensional reduction analyses using PCA and t-SNE.

We next tested whether macaque DE genes with expression that was significantly positively correlated (up-regulated DE genes) or significantly negatively correlated (down-regulated DE genes) with the measure of infection (*i.e.*, parasite load or parasitemia), infecting *Plasmodium* species, and the interaction between the measure of infection and the infecting *Plasmodium* species were enriched for particular biological functions. Specifically, we tested whether these genes were more likely than expected by chance to share molecular functions, biological processes, or cell component annotations from the Gene Ontology database^61, 62^. For each dataset, we also tested for functional enrichment of Human Phenotype Ontology^63^ annotations, using annotations for human orthologs matched to our macaque genes using the Ensembl gene mappings (accessed via g:Orth^64^). To test for enrichment, we used a hypergeometric test as implemented in g:Profiler^64^ and used all annotated genes after filtration for low read count as a comparative background list for differential expression determination. We finally tested whether these genes were more likely than expected by chance to also be differentially expressed in human response to *Plasmodium* parasites (*i.e.*, *P. vivax* and *P. falciparum*^34^). We did so using a hypergeometric test after converting our parasite species-DE genes to their one-to-one human orthologs in g:Profiler^64^.

### Identification of differentially expressed Plasmodium genes between malaria species

To identify *Plasmodium* genes with expression correlated to parasite load, we analyzed the combined datasets (PRJNA400695 and PRJNA388645) after collapsing *Plasmodium* genes into orthologous groups based on on one-to-one orthologs between *P. coatneyi* and *P. cynomolgi* (Tables S5-S6), which were gathered from PlasmoDB^56^. Using the combined read count matrix, we first calculated the normalization factors and, because there were very few *Plasmodium* reads, filtered for lowly expressed genes using the filterByExpr function of edgeR which considers datasets relative to their design matrix^58^. Using these calculated normalization factors, we then used the limma-voom package^59, 60^ to include biological replicates as a random effect (*i.e.*, blocking variable) and parasite load as a fixed effect in the following model:

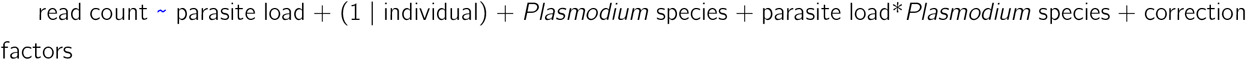

where, as above, correction factors include those adjusting for differences in number of reads and library complexity. We corrected for multiple tests via FDR and again considered a gene to be a parasitemia-DE gene if the adjusted *p*-value was less than 0.05. We then used Empirical Bayes *t*-statistics to identify significantly up- and down-regulated parasitemia-response genes.

Finally, to investigate potential biological relationships between parasite load and *Plasmodium* species identity using normalized and filtered read counts, we performed dimensional reduction analyses including PCA and t-SNE analyses.

Similar to the method used for macaque genes, we next tested whether *P. coatneyi-P. cynomolgi* orthologous gene pairs with expression that was significantly positively correlated (up-regulated) or significantly negatively correlated (down-regulated) with the measure of infection (*i.e.*, parasite load or parasitemia), *Plasmodium* species, or the interaction between the measure of infection and the *Plasmodium* species were enriched for particular functional annotations using Gene Ontology analysis as implemented in PlasmoDB^56^.

Code used in analyses is available at https://github.com/ambertrujillo/Comparative_macaque.

## Acknowledgements

We thank the MaHPIC researchers for making their data available. Funding provided by the Ford Foundation and N.S.F. BCS-2118108 to A.E.T.

## Author contributions statement

A.E.T. conceived the study, analyzed the results, and wrote the manuscript. C.M.B. conceived the study, interpreted the results, and wrote the manuscript. All authors reviewed the manuscript.

## Data availability statement

Data used in the present study are available from the National Center for Biotechnology Information (NCBI) Sequence Read Archive (SRA) under accessions PRJNA400695 and PRJNA388645.

## Additional information

The authors declare no competing interests.

